# Inhibition of peroxisomal protein PRX-11 promotes longevity in *Caenorhabditis elegans* via enhancements to mitochondria

**DOI:** 10.1101/2025.05.28.656437

**Authors:** Yash Flora, Dhriti Shastri, K. Adam Bohnert

## Abstract

Peroxisomes execute essential functions in cells, including detoxification and lipid oxidation. Despite their centrality to cell biology, the relevance of peroxisomes to aging remains understudied. We recently reported that peroxisomes are degraded *en masse* via pexophagy during early aging in the nematode *Caenorhabditis elegans*, and we found that downregulating the peroxisome-fission protein PRX-11/PEX11 prevents this age-dependent pexophagy and extends lifespan. Here, we further investigated how *prx-11* inhibition promotes longevity. Remarkably, we found that reducing peroxisome degradation with age led to concurrent improvements in another organelle: mitochondria. Animals lacking *prx-11* function showed tubular, youthful mitochondria in older ages, and these enhancements required multiple factors involved in mitochondrial tubulation and biogenesis, including FZO-1/Mitofusin, UNC-43 protein kinase, and DAF- 16/FOXO. Importantly, mutation of each of these factors negated lifespan extension in *prx-11-*defective animals, indicating that pexophagy inhibition promotes longevity only if mitochondrial health is co-maintained. Our data support a model in which peroxisomes and mitochondria track together with age and interdependently influence animal lifespan.

## Introduction

Age-related decline in bodily function is associated with increased disease incidence and higher mortality rates [1, 2]. Though the effects of aging are perhaps most immediately recognizable at a physiological level, health decline in older animals is rooted in dysfunction at the level of cells and molecules [3, 4]. Understanding the cellular changes that instigate age-related deterioration might suggest ways to curb their progression and promote longer, healthier lives.

Research over the past few decades has revealed a growing list of cellular hallmarks of aging that characterize an aged state [4]. These cellular aging hallmarks include mitochondrial stress, protein aggregation, telomere shortening, and deregulated signaling [4-8]. Using the nematode *Caenorhabditis elegans*, we recently identified another cellular change common to older animals: loss of peroxisomes via autophagic destruction [9]. Although peroxisomes have received comparably less attention than other organelles in the aging field, peroxisomes are increasingly recognized as essential modulators of cellular-stress responses and metabolic coordination. Indeed, their resident enzyme, catalase, breaks down hydrogen peroxide, and they contain additional enzymes that direct lipid oxidation in concert with mitochondria [10-12]. In our previous study, we developed a tandem-fluorophore reporter to track peroxisome degradation through autophagy (“pexophagy”) during *C. elegans* aging [9]. Whereas wild-type *C. elegans* showed a dramatic reduction in peroxisome number by day 5 of adulthood, we found that inhibiting a single peroxisome protein, PEX11 homolog PRX-11, prevented early-age pexophagy [9], most likely by interfering with the normal role of PRX-11/PEX11 in peroxisome fission [13]. Notably, *prx-11*-defective animals not only retained peroxisomes later in life than control animals, but they lived longer, too [9]. These findings support a model in which early-age autophagic destruction of peroxisomes might ultimately limit animal longevity. However, why pexophagy inhibition increases animal lifespan is poorly understood.

An important consideration in geroscience research is that aging animals are aging systems, and individual aging hallmarks often do not change in isolation. Due to interconnectivities in cellular architecture and signaling, changes to one hallmark can influence others. Among the other parts of the cell, mitochondria show the most obvious connections to peroxisomes. Peroxisomes and mitochondria are metabolically coupled through shared lipid substrates and reactive oxygen species (ROS) signaling, and both organelles rely on common fission machinery [14-17]. Despite these connections, it is unknown whether changes to peroxisomes during aging would have any effect on mitochondria. In many organisms, mitochondrial networks naturally undergo fragmentation with age [18-20]. In *C. elegans*, this age-associated mitochondrial fragmentation can be easily detected in body wall muscle, where tubular mitochondrial networks present in early adulthood progressively collapse into punctate, fragmented structures by mid-age [20, 21]. These changes to mitochondrial morphology are accompanied by mitochondrial membrane depolarization, mitochondrial calcium (Ca^2+^) accumulation, and locomotory decline [21-23]. Although mitochondrial quality-control pathways comprising fusion/fission machinery, mitophagy factors, and stress-responsive transcriptional programs can help to maintain the health of this organelle [24-27], the initiating events that destabilize mitochondrial homeostasis during early aging remain unclear. In principle, this might be linked to collapse in functionality of related organelles, including peroxisomes. If so, boosting peroxisomes in older animals might help to maintain more youthful mitochondria. Though an attractive hypothesis, this has yet to be experimentally tested.

In this study, we investigated whether enhanced longevity upon pexophagy inhibition involves coordinated changes to mitochondria. We found that *prx-11* downregulation enhanced multiple indications of mitochondrial health with age; namely, inhibiting *prx-11* function preserved mitochondrial network architecture, suppressed mitochondrial Ca^2+^ accumulation, and reduced oxidative stress in older animals. Moreover, these changes were associated with not only lifespan extension but also improvements to locomotory function, indicating that inhibition of peroxisome degradation protects mitochondrial integrity and delays associated aspects of physiological aging. We also found that the mitochondrial improvements observed in *prx-11*-defective animals required the mitochondrial fusion factor FZO-1 [27-29], the UNC-43 protein kinase, which opposes mitochondrial fission [30], and DAF-16, the *C. elegans* FOXO transcription factor [31], suggesting that mitochondrial-remodeling and biogenesis pathways are engaged downstream of peroxisome retention to promote mitochondrial health in older animals. Blocking the function of these factors also prevented *prx-11* knockdown from extending lifespan. Thus, we conclude that mitochondria are an integral component of the pro-longevity response triggered by pexophagy inhibition, and we posit that the rate of age-related peroxisome loss influences lifespan in part by affecting this separate organelle. This work establishes a new framework for understanding how inter-organelle crosstalk contributes to aging and highlights peroxisome turnover as a potential target for interventions aimed at preserving mitochondrial function and extending lifespan.

## Results

### Animals lacking *prx-11* function retain tubular mitochondria in older ages

We began our investigations by probing whether the enhanced longevity caused by *prx-11* RNA interference (RNAi) was associated with any other cellular changes indicative of slower aging. Because peroxisomes and mitochondria share some functional overlap [32, 33], we reasoned that mitochondria might be altered when pexophagy was inhibited. To test this, we visualized mitochondrial morphology using a mitochondrially targeted green fluorescent protein (Mito-GFP) expressed in *C. elegans* muscles [34]. We confirmed that animals treated with control RNAi showed typical age-dependent mitochondrial fragmentation in mid-and late-adulthood (Figure 1A), as indicated by decreased mitochondrial lengths and decreased junctions per object (Figure 1B,C). Remarkably, this was not the case in *prx-11*-RNAi animals; at days 5 and 10 of adulthood, *prx-11*-RNAi animals retained very tubular mitochondria, which resembled those of young, day 2 adult animals (Figure 1A-C). In contrast, knockdown of HMG-CoA reductase *hmgr-1,* which accelerates age-dependent pexophagy [9], induced some mitochondrial fragmentation evident from day 2 of adulthood (Figure 1A-C). We concluded that inhibition of pexophagy, as occurs by *prx-11* knockdown, is associated with retention of tubular, youthful mitochondria in older ages.

**Figure 1.**
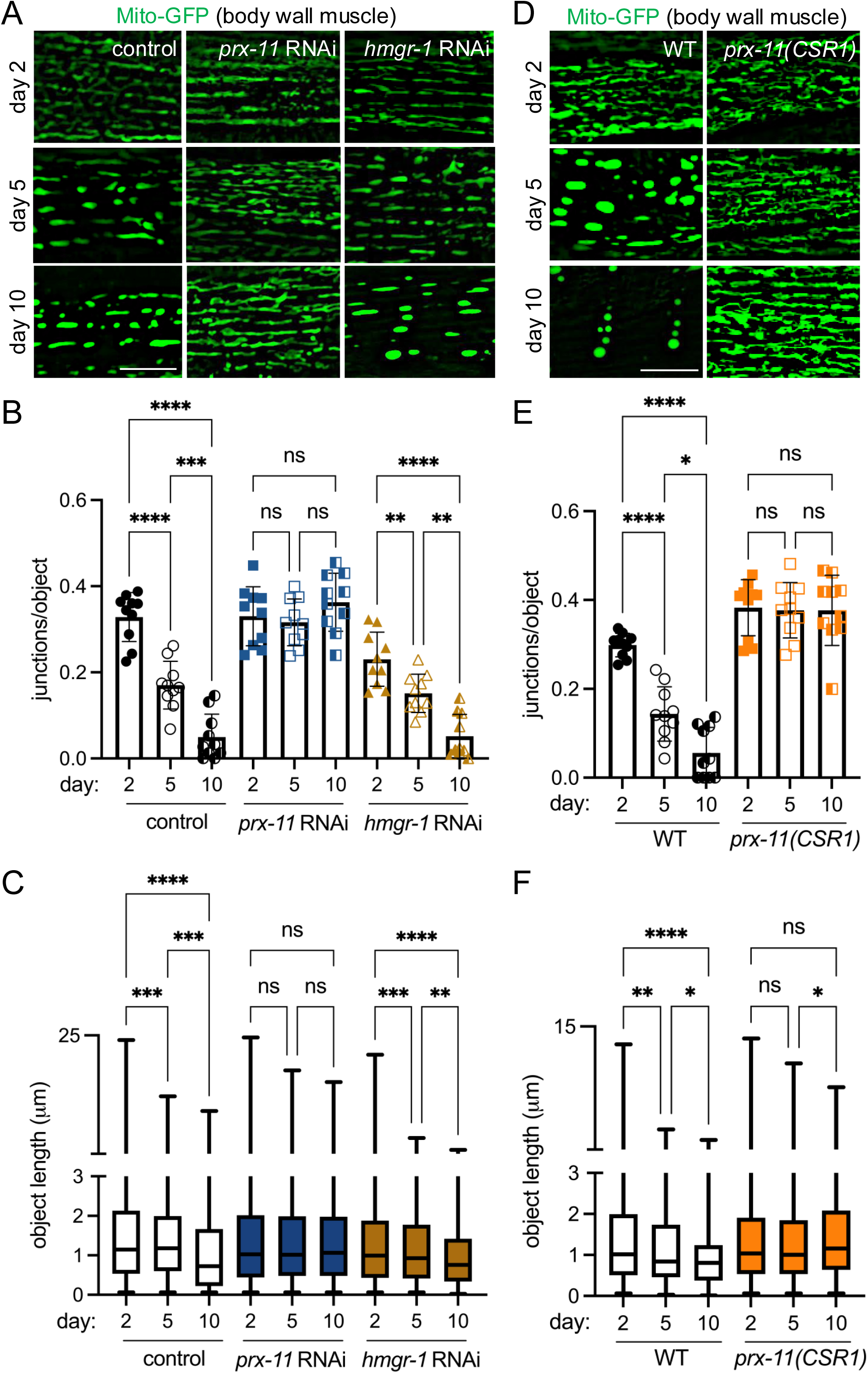
*prx-11* depletion maintains mitochondrial junctions and network morphology in aging *C. elegans*. (A) Representative images of Mito-GFP in body wall muscle of day 2, day 5, and day 10 adult hermaphrodite animals fed either control, *prx-11,* or *hmgr-1* RNAi. (B) Quantification of mitochondrial junctions per object in day 2, day 5, and day 10 adult hermaphrodite animals fed either control, *prx-11,* or *hmgr-1* RNAi (*n* = 10 worms per condition). Data are presented as mean ± SD. ****, *p ≤* 0.0001; ***, *p ≤* 0.001; **, *p ≤* 0.01; ns, not significant. One-way ANOVA with Tukey’s multiple comparisons. (C) Mito-GFP object lengths in day 2, day 5, and day 10 adult hermaphrodite animals fed either control, *prx-11,* or *hmgr-1* RNAi (*n* = 10 worms per condition). Data are presented as box-and-whisker plots (minimum, 25^th^ percentile, median, 75^th^ percentile, maximum). ****, *p ≤* 0.0001; ***, *p ≤* 0.001; **, *p ≤* 0.01; ns, not significant. One-way ANOVA with Tukey’s multiple comparisons. (D) Representative images of Mito-GFP in body wall muscle of day 2, day 5, and day 10 adult wild-type and *prx-11(CSR1)* hermaphrodite animals. (E) Quantification of mitochondrial junctions per object in day 2, day 5, and day 10 adult wild-type and *prx-11(CSR1)* hermaphrodite animals (*n* = 10 worms per condition). Data are presented as mean ± SD. ****, *p ≤* 0.0001; *, *p ≤* 0.05; ns, not significant. One-way ANOVA with Tukey’s multiple comparisons. (F) Mito-GFP object lengths in day 2, day 5, and day 10 adult wild-type and *prx-11(CSR1)* hermaphrodite animals (*n* = 10 worms per condition). Data are presented as box-and-whisker plots (minimum, 25^th^ percentile, median, 75^th^ percentile, maximum). ****, *p ≤* 0.0001; **, *p ≤* 0.01; *, *p ≤* 0.05; ns, not significant. One-way ANOVA with Tukey’s multiple comparisons. Bars: 10 μm.

To corroborate this result, we tested whether the same would be true in a previously generated *prx-11* mutant, *prx-11(CSR1)*, which contains an internal genetic deletion [35]. First, we verified that *prx-11(CSR1)* mutants, like *prx-11*-RNAi animals [9], show reduced age-dependent pexophagy (Supplementary Figure 1A,B), larger peroxisomes (Supplementary Figure 1C,D), and increased lifespans (Supplementary Figure 1E). Though loss of *prx-11* slowed pexophagy, we noted that it did not appear to impede bulk autophagy, using a tandemly tagged SQST-1/p62 autophagy receptor marker [36] (Supplementary Figure 1F,G). Importantly, *prx-11(CSR1)* mutant animals exhibited robust mitochondrial tubulation in older ages (Figures 1D-F), confirming that enhanced mitochondrial tubulation is a general result of *prx-11* loss of function. Thus, inhibiting *prx-11* not only leads to peroxisome retention but also preserves mitochondria with morphological characteristics of a more youthful state.

### Aged *prx-11*-RNAi animals show reductions in a second mitochondrial hallmark of aging, mitochondrial Ca^2+^

We were curious whether animals having loss of function in *prx-11* would display other molecular characteristics of youthful mitochondria in addition to enhanced tubulation. An increase in mitochondrial Ca^2+^ uptake has been observed with age in both worms and mice and has been associated with age-related diseases such as sarcopenia [22, 37] . Interestingly, blocking mitochondrial Ca^2+^ uptake via inhibition of the Ca^2+^ uniporter *mcu-1* [38, 39] increases mitochondrial tubulation in older animals [22], though this can perhaps be even further enhanced by *prx-11* knockdown (Supplementary Figure 2A,B). Given these considerations, we asked whether *prx-11*-RNAi animals showed reduced mitochondrial Ca^2+^ uptake in older ages. We used a mitochondrially targeted, low-affinity red fluorescent genetically encoded Ca^2+^ indicator for optical imaging (Mito-LAR-GECO) [22, 40] to test this hypothesis. Like *mcu-1* knockdown (Supplementary Figure 2A-C), *prx-11* knockdown caused diminished Mito-LAR-GECO signal in older animals, which trended together with improvements to mitochondrial morphology (Figure 2A-C). In contrast, accelerated pexophagy driven by *hmgr-1* knockdown was associated with premature Mito-LAR-GECO detection in young adulthood (Figure 2A,C). These data support the conclusion that retaining peroxisomes in older age contributes to maintenance of more youthful mitochondria.

**Figure 2.**
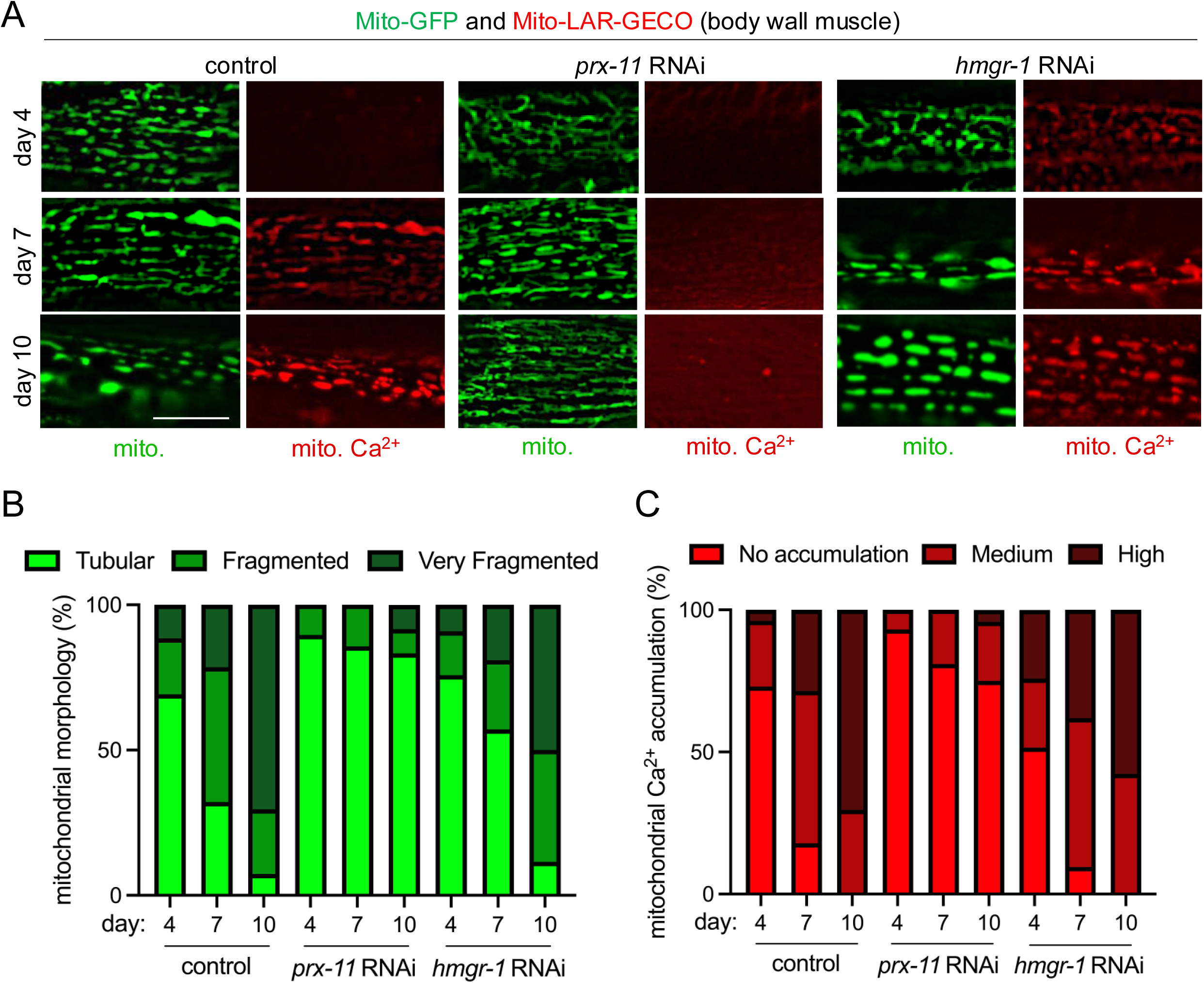
Enhanced mitochondrial tubulation in older *prx-*11-RNAi animals is associated with reduced mitochondrial Ca^2+^ accumulation. (A) Representative images of Mito-GFP and the Mito-LAR-GECO Ca^2+^ reporter in body wall muscle of day 4, day 7, and day 10 adult hermaphrodite animals fed either control, *prx-11,* or *hmgr-1* RNAi. (B,C) Classification of mitochondrial morphology and mitochondrial Ca^2+^ accumulation in day 4, day 7, and day 10 adult hermaphrodite animals fed either control, *prx-11,* or *hmgr-1* RNAi (*n* = 20 worms per condition). Bar: 10 μm.

### Loss of *prx-11* activity improves aspects of physiology related to mitochondrial health

We aimed to assess whether changes to physiology associated with altered mitochondrial function could be detected upon *prx-11* loss of function. We first evaluated whether *prx-11* knockdown had any effect on the mitochondrial unfolded protein response (UPR^mt^). UPR^mt^ is a stress-response pathway that, when activated, can improve the robustness of mitochondrial form and function and support increased longevity [41-43]. Though mitochondria appeared enhanced by *prx-11* knockdown, we did not observe a relative induction of UPR^mt^ in *prx-11*-RNAi animals, as judged using the UPR^mt^ reporters *Phsp-6*::GFP [43] and *Phsp-60*::GFP [43] (Supplementary Figure 3A-D). We also did not observe increased relative expression of the endoplasmic reticulum unfolded protein response (UPR^er^) reporter *Phsp-4*::GFP [44] in *prx-11*-RNAi animals (Supplementary Figure 3E,F), suggesting that organelle-based UPR pathways are generally not engaged by *prx-11* knockdown.

We next investigated whether *prx-11* knockdown was associated with changes in oxidative stress. A previous report linked loss of *prx-11* function to decreased oxidative stress [45]. However, this interpretation was made under the assumption that losing *prx-11* function would prevent peroxisome formation and lead to an absence of peroxisomes [45]. Our data indicate that losing *prx-11* function instead causes an increase in peroxisome area during animal aging (Supplementary Figure 1C,D) [9]. We therefore sought to reexamine this relationship. We found that *prx-11*-RNAi animals indeed exhibited a marked decrease in labeling by the ROS-sensitive dye dihydroethidium (DHE) at days 1 and 5 of adulthood (Figure 3A,B). Interestingly, this was associated with reduced expression of a superoxide dismutase transcriptional reporter, *Psod-3*::GFP [46] (Figure 3C,D). These data suggest that the reduction in ROS may not be due to increased superoxide-dismutase activity *per se* but instead may result from lower intrinsic ROS production, consistent with better mitochondrial function [47, 48].

**Figure 3.**
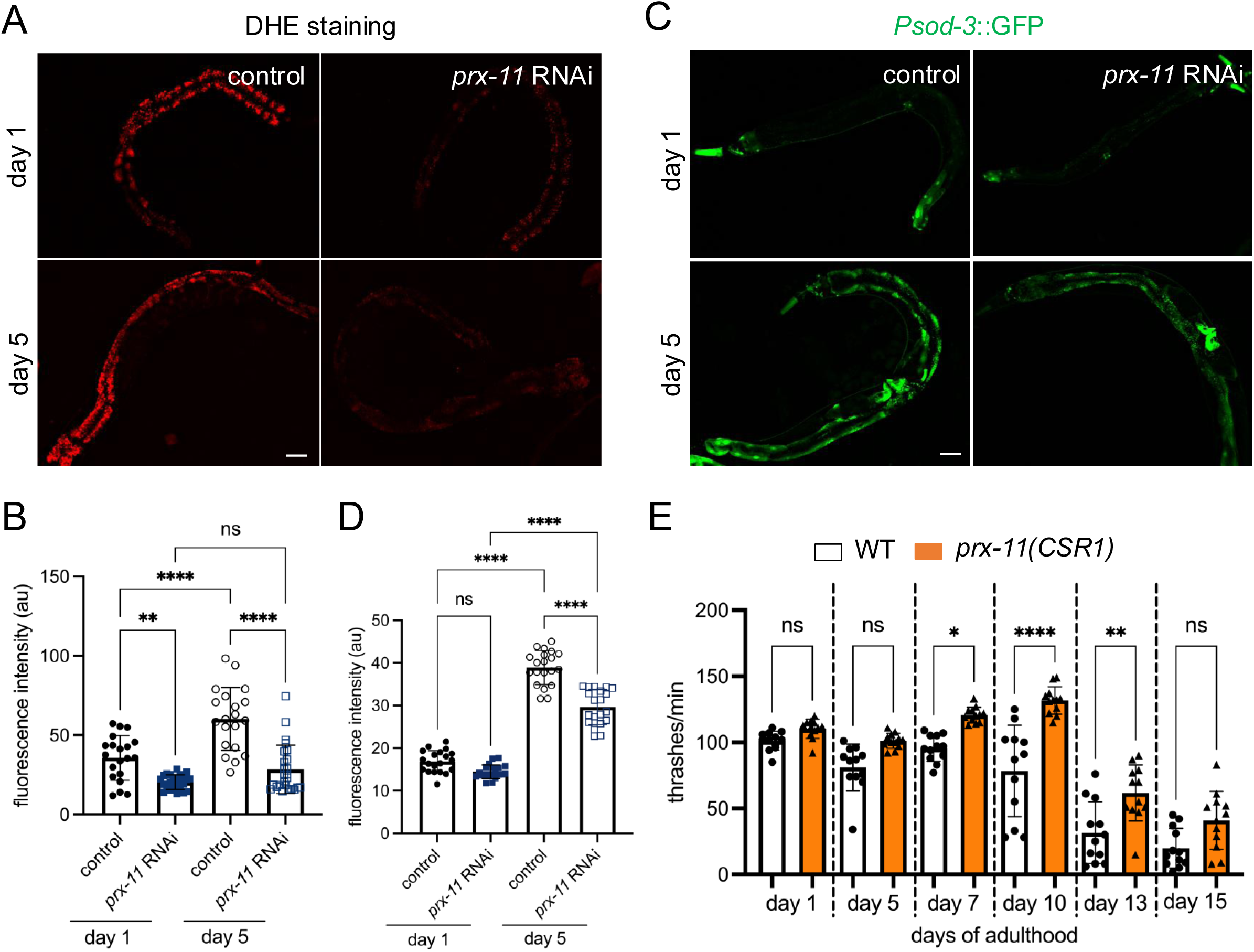
*prx-11* depletion attenuates oxidative stress and preserves motor function during aging. (A) Representative images of DHE-stained day 1 and day 5 adult hermaphrodite animals fed either control or *prx-11* RNAi. (B) Quantification of DHE staining fluorescence intensities in day 1 and day 5 adult hermaphrodite animals fed either control or *prx-11* RNAi (*n* = 20 worms per condition). Data are presented as mean ± SD. ****, *p ≤* 0.0001; **, *p ≤* 0.01; ns, not significant. One-way ANOVA with Tukey’s multiple comparisons. (C) Representative images of *Psod-3*::GFP in day 1 and day 5 adult hermaphrodite animals fed either control or *prx-11* RNAi. (D) Quantification of *Psod-*3::GFP fluorescence intensities in day 1 and day 5 adult hermaphrodite animals fed either control or *prx-11* RNAi (*n* = 20 worms per condition). Data are presented as mean ± SD. ****, *p ≤* 0.0001; ns, not significant. One-way ANOVA with Tukey’s multiple comparisons. (E) Quantification of thrashing rates in adult wild-type and *prx-11(CSR1)* hermaphrodite animals from day 1 through day 15 of adulthood (*n* = 12 worms per condition). Data are presented as mean ± SD. ****, *p ≤* 0.0001; **, *p ≤* 0.01; *, *p ≤* 0.05; ns, not significant. One-way ANOVA with Tukey’s multiple comparisons. Bars: 100 μm.

In *C. elegans*, preserved mitochondrial function with age is also associated with improved muscle performance [21, 36, 49]. This can be assessed by measuring locomotory activity, which in *C. elegans* is evaluated by counting the number of body thrashes in liquid medium per unit time [36, 50]. We found that loss of *prx-11* function in fact enhanced locomotory activity of aging *C. elegans* (Figure 3E). These data are also consistent with improved muscle mitochondrial function and indicate that *prx-11* loss of function not only extends lifespan but boosts health in older animals.

### Mitochondrial tubulation in aged *prx-11*-RNAi animals requires FZO-1/Mitofusin, UNC-43 protein kinase, and DAF-16/FOXO

Given the longevity and physiological benefits associated with *prx-11* knockdown, we sought to identify genes required for enhanced mitochondrial tubulation in older animals under this condition. A likely candidate was *fzo-1*, which encodes a Mitofusin GTPase essential for mitochondrial fusion [28, 29, 51]. We found that *fzo-1* mutant animals showed hyper-fragmented mitochondria, even when *prx-11* was knocked down (Figure 4A-C). Thus, FZO-1/Mitofusin activity is required for enhanced mitochondrial tubulation upon *prx-11* knockdown. Typically, dynamin-related DRP-1, which promotes mitochondrial fission, counteracts the activity of FZO-1/Mitofusin [29, 51, 52]. DRP-1 itself is inhibited by UNC-43, a kinase that blocks DRP-1 activity via phosphorylation [30]. We found that mutation or knockdown of *unc-43*, like mutation of *fzo-1*, dampened mitochondrial tubulation in older *prx-11*-RNAi animals (Figure 4A-C and Supplementary Figure 4A-C). Thus, key proteins directly involved in the mitochondrial fusion/fission cycle feed into the regulation of mitochondrial biology in this genetic background.

**Figure 4.**
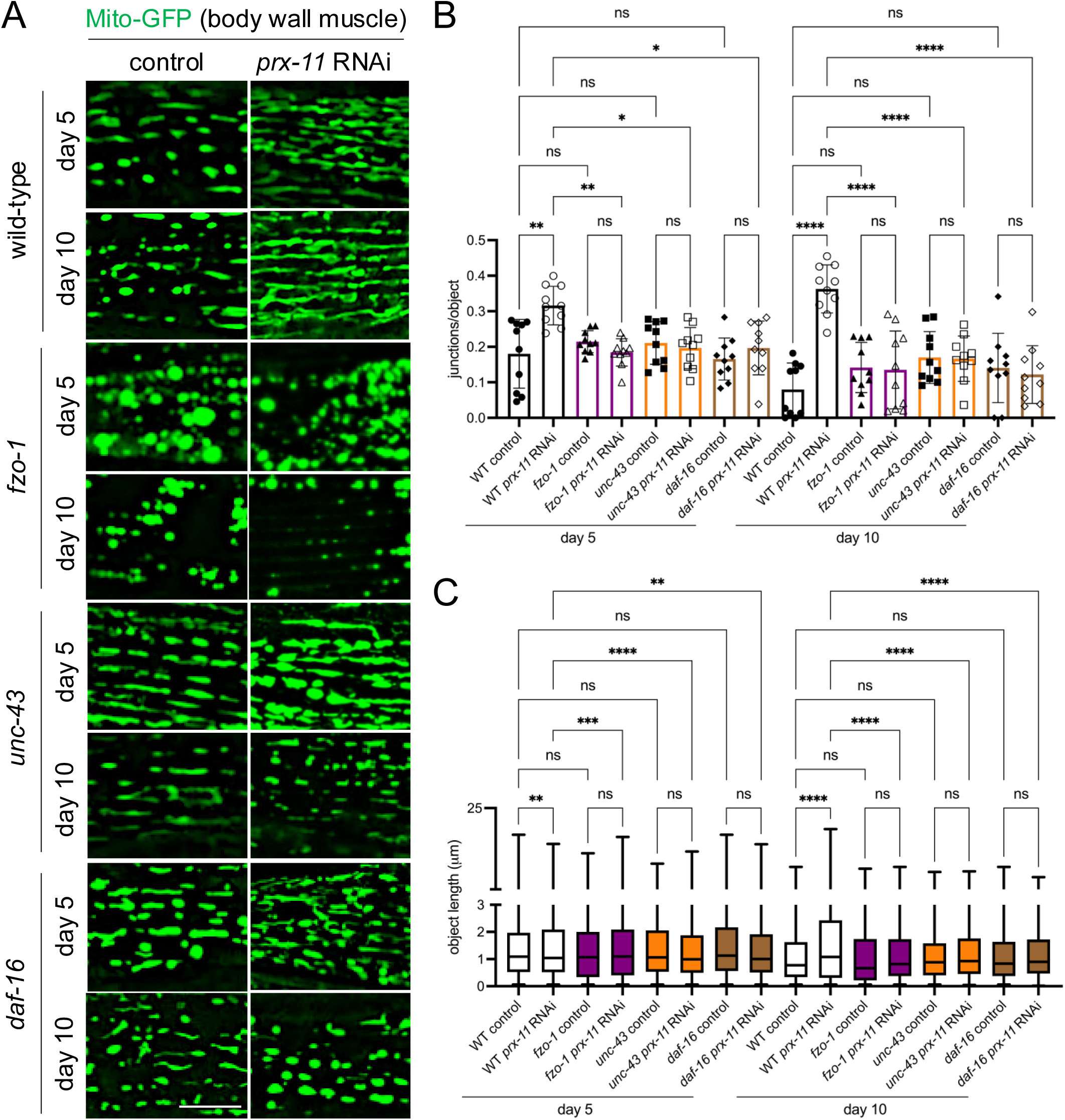
Loss of *fzo-1, unc-43*, or *daf-16* abrogates mitochondrial network preservation in *prx-11* knockdown animals. (A) Representative images of Mito-GFP in body wall muscle of day 5 and day 10 adult wild-type, *fzo-1, unc-43*, and *daf-16* hermaphrodite animals fed either control or *prx-11* RNAi. (B) Quantification of mitochondrial junctions per object in day 5 and day 10 adult wild-type, *fzo-1*, *unc-43*, and *daf-16* hermaphrodite animals fed either control or *prx-11* RNAi (*n* = 10 worms per condition). Data are presented as mean ± SD. ****, *p ≤* 0.0001; **, *p ≤* 0.01; *, *p ≤* 0.05; ns, not significant. One-way ANOVA with Tukey’s multiple comparisons. (C) Mito-GFP object lengths in day 5 and day 10 adult wild-type, *fzo-1*, *unc-43*, and *daf-16* hermaphrodite animals fed either control or *prx-11* RNAi (*n* = 10 worms per condition). Data are presented a box-and-whisker plots (minimum, 25^th^ percentile, median, 75^th^ percentile, maximum). ****, *p ≤* 0.0001; ***, *p ≤* 0.001; **, *p ≤* 0.01; ns, not significant. One-way ANOVA with Tukey’s multiple comparisons. Bar: 10 μm.

We also tested whether the improvements to mitochondria seen upon *prx-11* knockdown required function of DAF-16/FOXO, a transcription factor that acts as a master regulator of longevity regulation and mitochondrial homeostasis [27, 31, 53]. Indeed, mutating or knocking down *daf-16* reduced mitochondrial tubulation in older *prx-11*-RNAi animals (Figure 4A-C and Supplementary Figure 4A-C). These results identify several factors involved in mitochondrial regulation and biogenesis that are important for retention of tubular mitochondria with age upon loss of *prx-11* function.

### Preventing mitochondrial tubulation negates lifespan extension in *prx-11*-RNAi animals

We next asked whether mitochondrial enhancements were causal in lifespan extension upon *prx-11* RNAi. We performed lifespan analysis of wild-type, *fzo-1*, *unc-43*, and *daf-16* animals on control and *prx-11* RNAi. Though *prx-11* RNAi reproducibly extended the lifespan of wild-type animals as previously reported [9, 45], it failed to extend lifespan in any of the three mutant backgrounds where *prx-11* RNAi was insufficient to drive mitochondrial tubulation in older age (i.e., *fzo-1, unc-43*, and *daf-16*) (Figure 5A-C). We concluded that *prx-11*-RNAi animals require improvements to mitochondria in order to live longer. Thus, one mechanism contributing to enhanced lifespan upon pexophagy inhibition is a coordinated improvement to mitochondrial biology.

**Figure 5.**
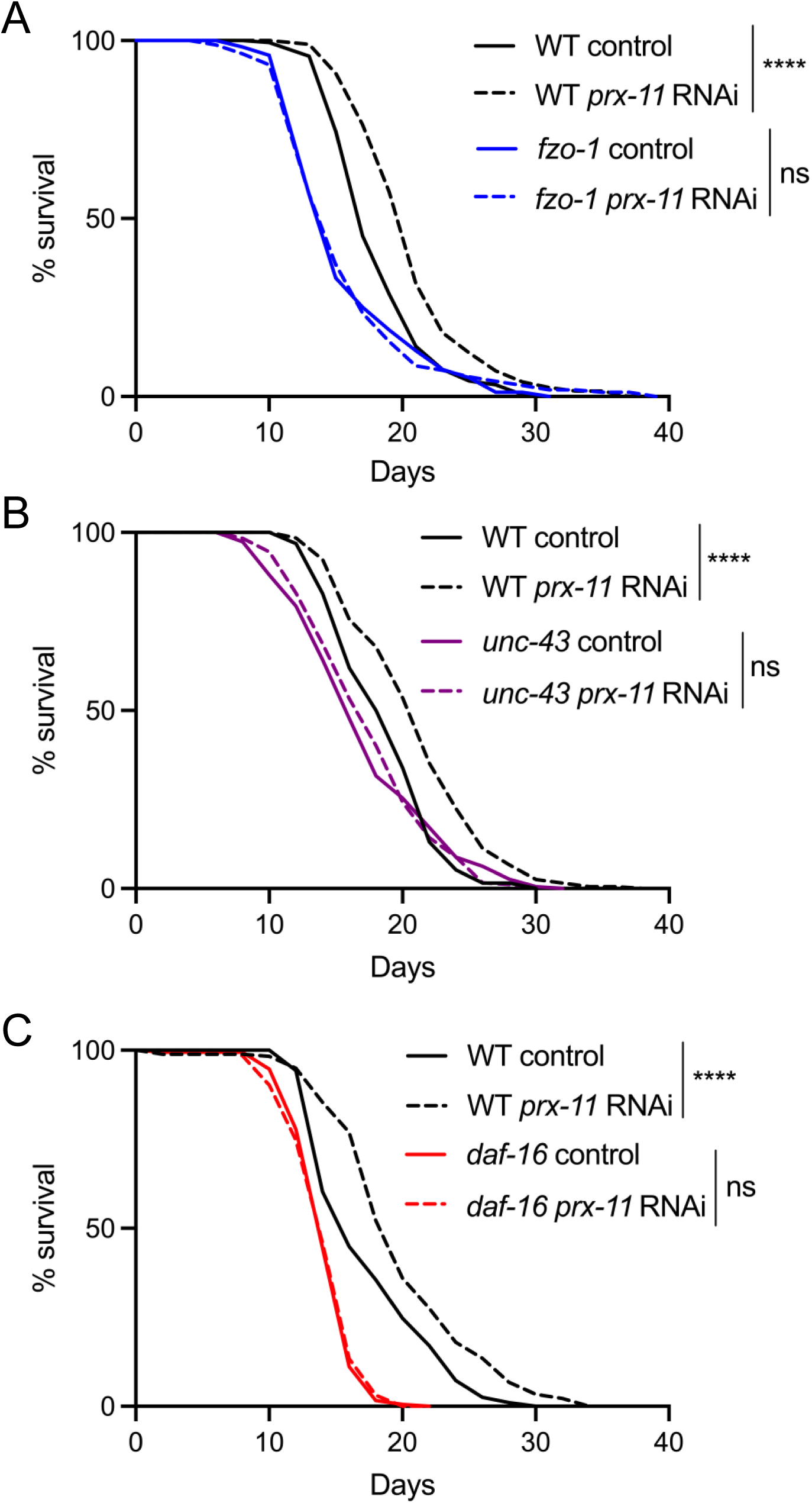
*fzo-1, unc-43*, and *daf-16* mutations block lifespan extension upon *prx-11* knockdown. (A) Lifespan analysis of wild-type and *fzo-1* hermaphrodite animals fed either control or *prx-11* RNAi. Significance was determined using a log-rank test. Wild-type control v. wild-type *prx-11* RNAi: Χ^2^= 34.61, *p ≤* 0.0001; *fzo-1* control v. *fzo-1 prx-11* RNAi: Χ^2^ = 0.01, ns, not significant. (B) Lifespan analysis of wild-type and *unc-43* hermaphrodite animals fed either control or *prx-11* RNAi. Significance was determined using a log-rank test. Wild-type control v. wild-type *prx-11* RNAi: Χ^2^ = 31.86, *p ≤* 0.0001; *unc-43* control v. *unc-43 prx-11* RNAi: Χ^2^ = 0.03, ns, not significant. (C) Lifespan analysis of wild-type and *daf-16* hermaphrodite animals fed either control or *prx-11* RNAi. Significance was determined using a log-rank test. Wild-type control v. wild-type *prx-11* RNAi: Χ^2^ = 24.21, *p ≤* 0.0001; *daf-16* control v. *daf-16 prx-11* RNAi: Χ^2^ = 0.13, ns, not significant.

## Discussion

Our findings identify peroxisome degradation as a modifiable event that controls mitochondrial aging (Figure 6). This provides a framework for understanding how inter-organelle communication sculpts the aging trajectory and suggests that targeting peroxisome homeostasis may offer a novel approach to preserve cellular function and extend healthy lifespan (Figure 6). These data expand on our previous work, which demonstrated that peroxisome loss begins early in adulthood [9], before overt signs of mitochondrial stress or damage are detectable. While mitochondrial quality control has traditionally been studied in isolation, our data support a model in which early degradation of peroxisomes acts as a gatekeeper event that modulates downstream changes to mitochondria.

**Figure 6.**
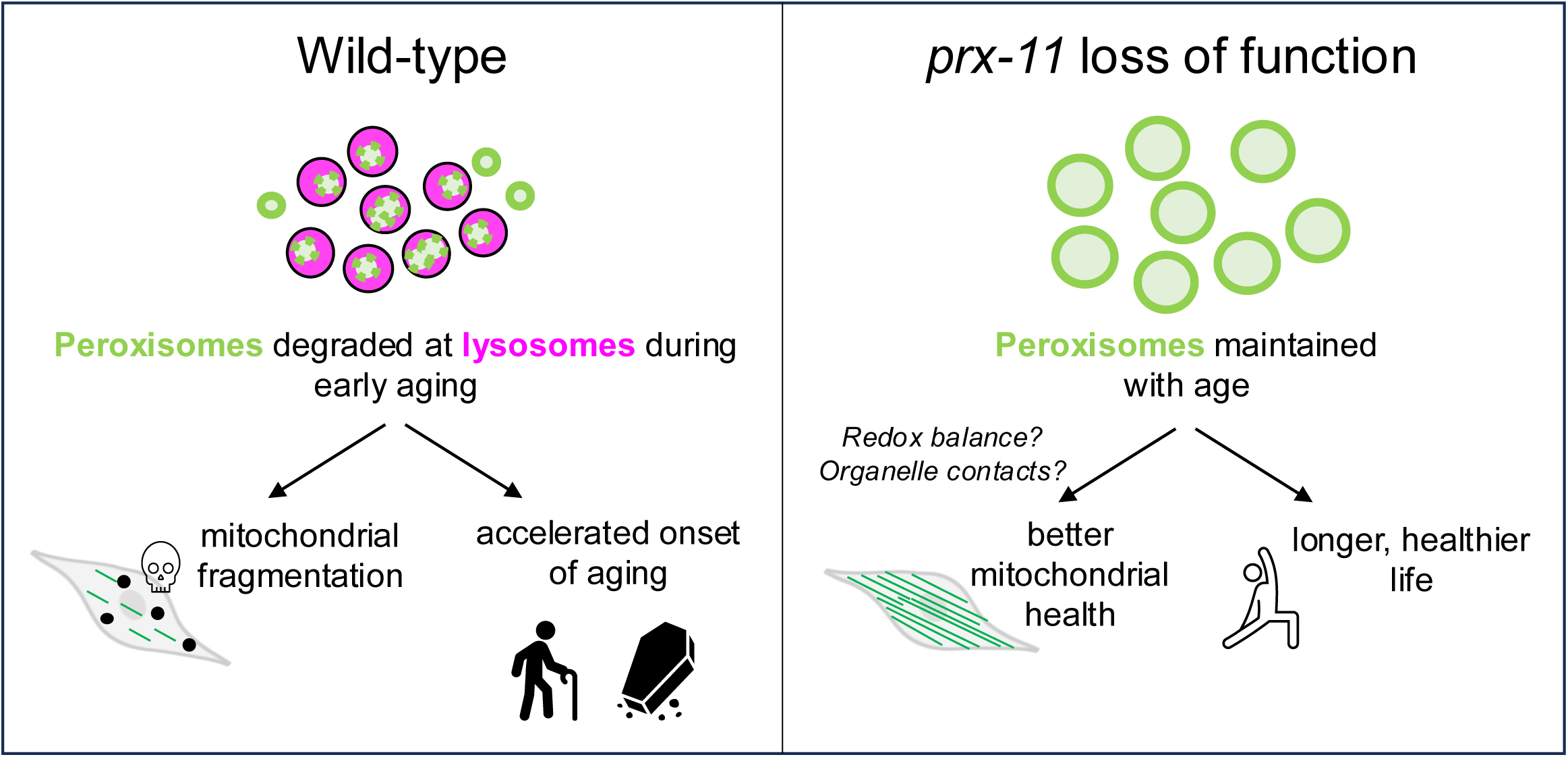
Model for the role of PRX-11 in regulating peroxisome degradation and mitochondrial homeostasis during aging. The schematic in the left panel illustrates that PRX-11 promotes age-associated peroxisome degradation and, in doing so, likely contributes to mitochondrial fragmentation and functional decline in older wild-type animals. The schematic in the right panel illustrates that loss of PRX-11 activity, via gene knockdown or mutation, inhibits pexophagy and preserves peroxisome abundance in older animals, thereby supporting enhanced mitochondrial integrity and lifespan extension. The effects of peroxisome retention on mitochondria quality with age involve crosstalk between the two organelles, potentially mediated by changes to physical interactions or metabolic signaling.

The nature of the peroxisomes retained upon loss of *prx-11* function is somewhat mysterious. In both *prx-11*-RNAi and mutant animals, peroxisomes not only persist with age but also increase in size. It is unclear whether the observed increase in peroxisome size enables them to evade degradation due to impaired fission or altered lysosomal targeting. Peroxisome size is known to influence degradation in other systems; larger or hyper-elongated peroxisomes have been shown to resist autophagic capture [54]. This raises the possibility that *prx-11* loss leads to a population of “autophagy-resistant” peroxisomes that remain structurally intact. Our data showing that SQST-1 turnover is unimpeded in *prx-11*-RNAi animals suggest that these peroxisomes do not accumulate as autophagy intermediates but may rather bypass the degradative pathway altogether.

An interesting observation is that inhibiting *prx-11* prevents mitochondrial fragmentation and delays aging phenotypes without activating canonical stress response pathways such as the UPR^mt^ or UPR^er^. Instead, our genetic data reveal that mitochondrial maintenance and lifespan extension mediated by *prx-11* RNAi require specific regulators of mitochondrial homeostasis, including FZO-1, UNC-43, and DAF-16. How loss of PRX-11, a protein specifically localized to peroxisomes [35], subsequently influences the function and regulation of these mitochondrial regulators is an open question.

One possibility is that peroxisome retention helps to maintain redox balance or buffering capacity, thereby impacting mitochondria (Figure 6). This possibility is consistent with observed reductions in ROS and calcium accumulation in *prx-11*-RNAi animals. Peroxisomes are known to modulate intracellular redox through catalase activity and fatty-acid oxidation, both of which are tightly linked to mitochondrial function [32, 55]. In mammalian cells, peroxisome-derived NADH and acetyl-CoA feed into mitochondrial metabolism, and catalase plays a key role in buffering hydrogen peroxide produced by both peroxisomal and mitochondrial activity, thereby limiting mitochondrial oxidative damage [56, 57]. The phenotypic suppression of oxidative stress and calcium overload suggests that the retained peroxisomes in *prx-11* mutants are not inert. Conversely, animals treated with *hmgr-1* RNAi, which accelerates pexophagy [9], exhibited increased mitochondrial calcium accumulation, and animals with mutations in *hmgr-1* have previously been reported to show other mitochondrial defects [58]. These data support a model in which early peroxisome loss compromises mitochondrial stability by removing buffering capacity or key metabolic intermediates.

Another possibility is that peroxisomes may engage in direct or indirect signaling with mitochondria to regulate network dynamics and transcriptional adaptation (Figure 6). The requirement for FZO-1 implies that mitochondrial fusion is an essential downstream effector of *prx-11* loss. That DAF-16, UNC-43, and mitochondrial biogenesis regulators are also required supports a model in which peroxisome retention activates a broader remodeling program involving transcriptional and signaling components. Although this program has yet to be fully described, plausible components include redox-sensitive transcriptional regulators, lipid-signaling intermediates, or even physical tethering complexes between peroxisomes and mitochondria [27, 31, 57, 59]. Recent work in mammalian cells and yeasts has identified peroxisome-mitochondria contact sites that facilitate lipid exchange and coordinate fission events [60-62]; it is exciting to speculate that similar structures may exist in *C. elegans*, potentially contributing to peroxisome-mitochondrial crosstalk during aging.

An additional point of note is that our findings challenge the prevailing view that organelle degradation is uniformly beneficial during aging. While mitophagy and other selective autophagy pathways are essential for damage clearance and can promote longevity [63], our findings suggest that premature destruction of functional peroxisomes may act as a destabilizing event that initiates mitochondrial fragmentation and health decline. In *C. elegans*, animals having loss-of-function mutations in the mTORC2 complex show hyperactive autophagy but live short [64]. Interestingly, this short lifespan has been linked to mitochondrial dysfunction [64]. It will be important to investigate whether pexophagy is also accelerated in this genetic background, perhaps explaining scenarios where animals show high autophagic capacity but also have decreased lifespan and impaired mitochondrial functionality.

## Materials and Methods

### *C. elegans* strain generation and maintenance

*C. elegans* strains used in this study are listed in Supplementary Table 1. The *prx-11* mutant strain, *prx-11(CSR1)*, which carries a CRISPR-engineered deletion [35], was graciously provided by the Huanhu Zhu Lab. For genetic crosses, transgenes expressing fluorescent proteins were tracked by stereomicroscopy, and gene deletions and mutations were verified by nested polymerase chain reactions (PCRs) and/or sequencing.

*C. elegans* were maintained on NGM agar (51.3 mM NaCl [ThermoFisher Scientific, S271-10], 0.25% peptone [ThermoFisher Scientific, BP1420-500], 1.7% agar [ThermoFisher Scientific, BP1423-500], 1 mM CaCl_2_ [G-Biosciences, R040], 1 mM MgSO_4_ [ThermoFisher Scientific, M65-500], 25 mM KPO_4_ [KH_2_PO_4_, ThermoFisher Scientific, P285-500; K_2_HPO_4_, ThermoFisher Scientific, P288-500], 12.9 μM cholesterol [Alfa Aesar, A11470], pH 6.0) seeded with *E. coli* OP50 bacteria at 20°C. Synchronous populations of animals were obtained by bleaching adult hermaphrodites; animals were vortexed in 1 mL bleaching solution (0.5 mM NaOH [ThermoFisher Scientific, S318-1], 20% bleach [Clorox, E624362]) for 5 min and then washed three times in M9 buffer (22 mM KH_2_PO_4_ [ThermoFisher Scientific, P285-500], 42 mM Na_2_HPO_4_ [ThermoFisher Scientific, S374-1], 85.5 mM NaCl [ThermoFisher Scientific, S271-10], 1 mM MgSO_4_ [ThermoFisher Scientific, M65-500], pH 7.0) before isolated eggs were then plated on OP50-seeded NGM. For aging experiments, adult animals were transferred daily to fresh OP50 and/or RNAi-seeded plates to separate adults from progeny.

For RNAi experiments, L4-stage animals were transferred onto RNAi plates (NGM with 100 ng/μl carbenicillin [ThermoFisher Scientific, BP26481] and 1 mM IPTG [ThermoFisher Scientific, 34060]) seeded with control or RNAi bacteria. The RNAi constructs were obtained from the Julie Ahringer collection [65]. An empty L4440 vector transformed into *E. coli* HT115 was used as a negative control. Clones were verified by DNA sequencing.

### Microscopy and image analysis

Four-percent agarose (ThermoFisher Scientific, BP164) gel pads dried with a Kimwipe (Kimtech) were placed on Gold Seal™ glass microscope slides. A small volume of 10 mM levamisole (Acros Organics, AC18787) was spotted on the agarose gel pad, after which worms were transferred to the levamisole drop and a glass cover slip (ThermoFisher Scientific, 12-540-B) was placed on top to complete the mounting. Live-animal fluorescence microscopy was performed using a Leica DMi8 THUNDER imager, equipped with 10*X* (NA 0.32), 40*X* (NA 1.30), and 100*X* (NA 1.40) objectives, a DFC9000 GTC camera, and GFP and Texas Red filter sets.

Mitochondrial networks were analyzed using “Skeleton” plugin in FIJI/ImageJ. Briefly, images were converted to binary 8-bit images and then to skeleton images. Skeleton images were then quantified using the “Analyze Skeleton” function. Numbers of objects, numbers of junctions, and object lengths were scored. An “object” is defined by the “Analyze Skeleton” function as a branch connecting two endpoints, an endpoint and junction, or two junctions. Junctions/object was used as a parameter to quantify network complexity and integrity.

For mitochondrial Ca^2+^ accumulation analysis, animals were categorized into discrete Ca^2+^ accumulation states (low, moderate, or high) based on manually defined thresholds.

For fluorescence intensity analysis, regions of interest were outlined using the freehand selection tool in FIJI/ImageJ, and mean intensity values were obtained using the “Analyze>Measure” function. For all fluorescence intensity measurements, 20% laser intensity, 300 ms exposure time, and 100% Fluorescence Intensity Manager settings were used.

Individual peroxisome areas were calculated by freehand tracing peroxisomes in FIJI/ImageJ and using the “Analyze>Measure” function.

ROS levels were measured using DHE staining. Synchronized animals were exposed to RNAi from L4 stage and aged until day 5 of adulthood. On days 1 and 5, animals were washed twice in M9 buffer, then incubated in 3 μM DHE (VWR, 76482-322) in M9 buffer for 45 minutes at room temperature in the dark. After incubation, animals were washed twice to remove excess dye and mounted on agarose pads for imaging. Red fluorescence was quantified in individual animals using FIJI/ImageJ, and mean intensity values were compared across conditions.

### Lifespan analysis

Synchronous populations of animals were transferred as late L4s to OP50 and/or RNAi seeded plates. The OP50 and/or RNAi plates were spotted with 5 mM FUdR (Acros Organics, 227605000) to prevent progeny production. 200 animals were analyzed per condition. Animals that exploded, bagged, or crawled off plates were censored during analysis. Lifespans were analyzed using OASIS 2 software [66], and statistical significance was assessed using a log-rank test.

### Thrashing assay

Synchronous populations of animals were transferred as late L4s to NGM plates seeded with OP50 bacteria. Worms were transferred to fresh plates every day to separate adults from their progeny. To score thrashing rates, individual worms were transferred into a drop of M9 buffer on an NGM plate, and the number of body thrashes were counted in a 1-minute period.

### Statistical analysis

Data were statistically analysed using GraphPad Prism software. For two-sample comparisons, an unpaired *t*-test was used to determine significance (α = 0.05). For comparisons of three or more samples, a one-way ANOVA with Tukey’s multiple comparisons was used to determine significance (α = 0.05). Statistical significance of lifespan data was determined using a log-rank test.

## Abbreviations

Ca^2+^: calcium
DHE: dihydroethidium
GFP: green fluorescent protein
Mito-GFP: mitochondrially targeted green fluorescent protein;
Mito-LAR-GECO: mitochondrially targeted low-affinity red fluorescent genetically encoded Ca^2+^ indicator for optical imaging;
PCR: polymerase chain reaction;
RNAi: RNA interference;
ROS: reactive oxygen species;
UPR^mt^: mitochondrial unfolded protein response;
UPR^er^: endoplasmic reticulum unfolded protein response.

## Author Contributions

Conceptualization: YF, KAB; Methodology: YF, DS; Formal Analysis: YF, DS; Resources: YF, KAB; Data Curation: YF, DS; Visualization: YF, DS; Funding acquisition: KAB; Project administration: KAB; Supervision: KAB; Writing – original draft: YF, KAB; Writing – review & editing: YF, DS, KAB.

## Acknowledgments

We thank members of the Bohnert lab and the Alyssa Johnson lab at LSU for comments on the study and critical reading of the manuscript. Some strains were provided by the *Caenorhabditis* Genetics Center (CGC), which is funded by the National Institutes of Health Office of Research Infrastructure Programs grant P40OD010440. We also thank Huanhu Zhu for generously providing the *prx-11(CSR1)* strain.

