## Supplementary Figures and Table for "Inhibition of peroxisomal protein PRX-11 promotes longevity in *Caenorhabditis elegans* via enhancements to mitochondria"

Supplementary Figure 1

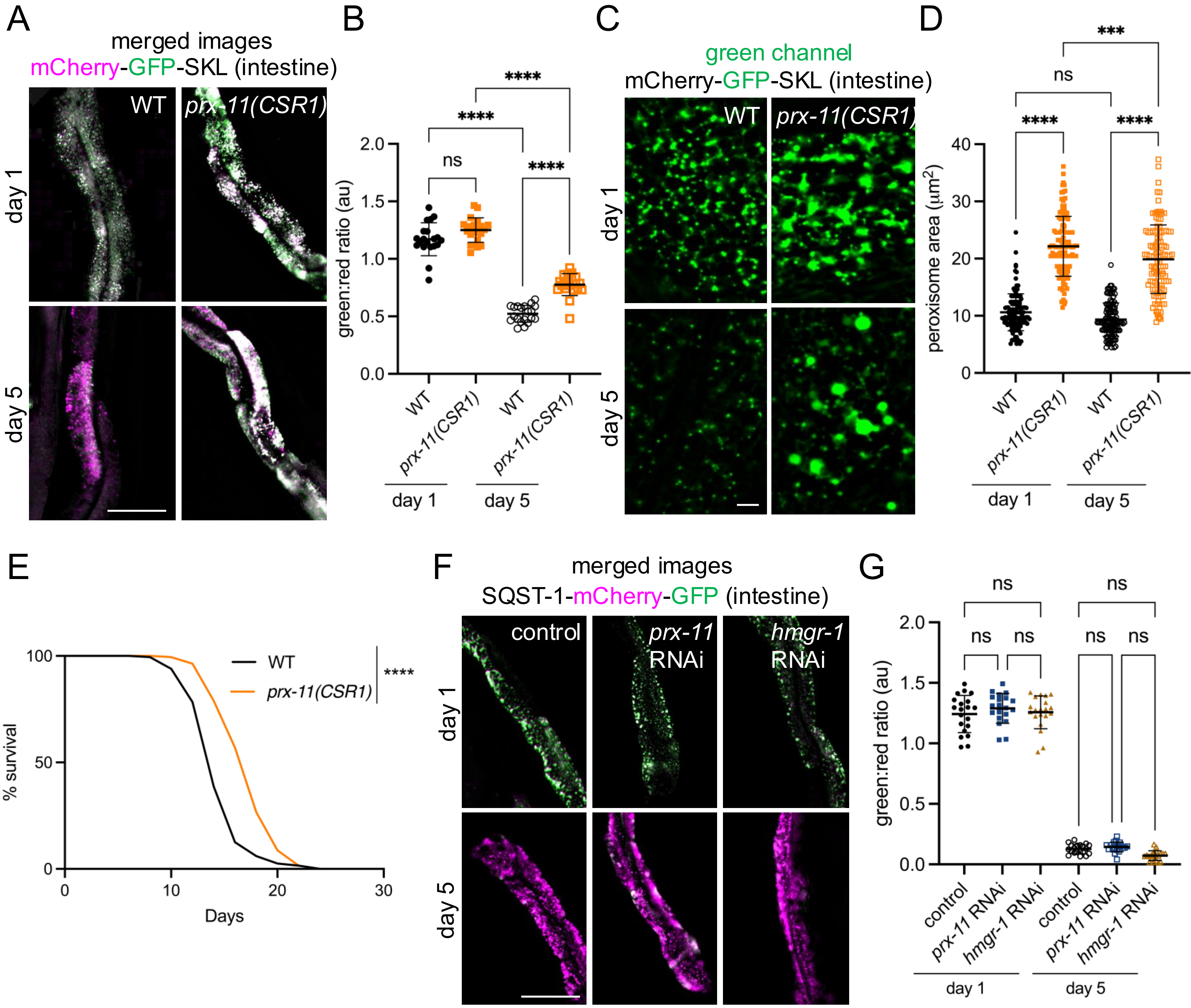

**Supplementary Figure 1. Loss of *prx-11* function, such as in *prx-11(CSR1)* mutants, is associated with peroxisome enlargement, lifespan extension, and a reduction in pexophagy but not autophagy in general.** (A) Representative merged green and red fluorescence images of the mCherry-GFP-SKL pexophagy reporter at day 1 and day 5 of adulthood in wild-type and *prx-11(CSR1)* animals. Pexophagy is indicated by a decrease in green fluorescence, which is sensitive to acidic pH, relative to red fluorescence, which is insensitive to acidic pH. Bar: 100 µm. (B) Quantification of green:red fluorescence ratios for mCherry-GFP-SKL at day 1 and day 5 of adulthood in wild-type and *prx-11(CSR1)* animals ($n$ = 20 worms per condition). Data are presented as mean ± SD. ****, $p \leq 0.0001$; ns, not significant. One-way ANOVA with Tukey's multiple comparisons. (C) Representative green channel images of mCherry-GFP-SKL at day 1 and day 5 of adulthood in wild-type and *prx-11(CSR1)* animals. Bar: 10 µm. (D) Quantification of individual peroxisome areas at day 1 and day 5 of adulthood in wild-type and *prx-11(CSR1)*. Data are presented as mean ± SD. ****, $p \leq 0.0001$; ***, $p \leq 0.001$; ns, not significant. One-way ANOVA with Tukey's multiple comparisons. (E) Lifespan analysis of wild-type and *prx-11(CSR1)* animals. Significance was determined using a log-rank test. Wild-type v. *prx-11(CSR1)*: $X^2 = 79.33$, $p \leq 0.0001$. (F) Representative merged green and red fluorescence images of the SQST-1-mCherry-GFP autophagy reporter at day 1 and day 5 of adulthood in animals fed either control, *prx-11,* or *hmgr-1* RNAi. Autophagic activity is indicated by a decrease in green fluorescence, which is sensitive to acidic pH, relative to red fluorescence, which is insensitive to acidic pH. Bar: 100 µm. (G) Quantification of green:red fluorescence ratios for SQST-1-mCherry-GFP at day 1 and day 5 of adulthood in animals fed either control, *prx-11,* or *hmgr-1* RNAi (*n* = 20 worms per condition). Data are presented as mean ± SD. ns, not significant. One-way ANOVA with Tukey’s multiple comparisons.

Supplementary Figure 2

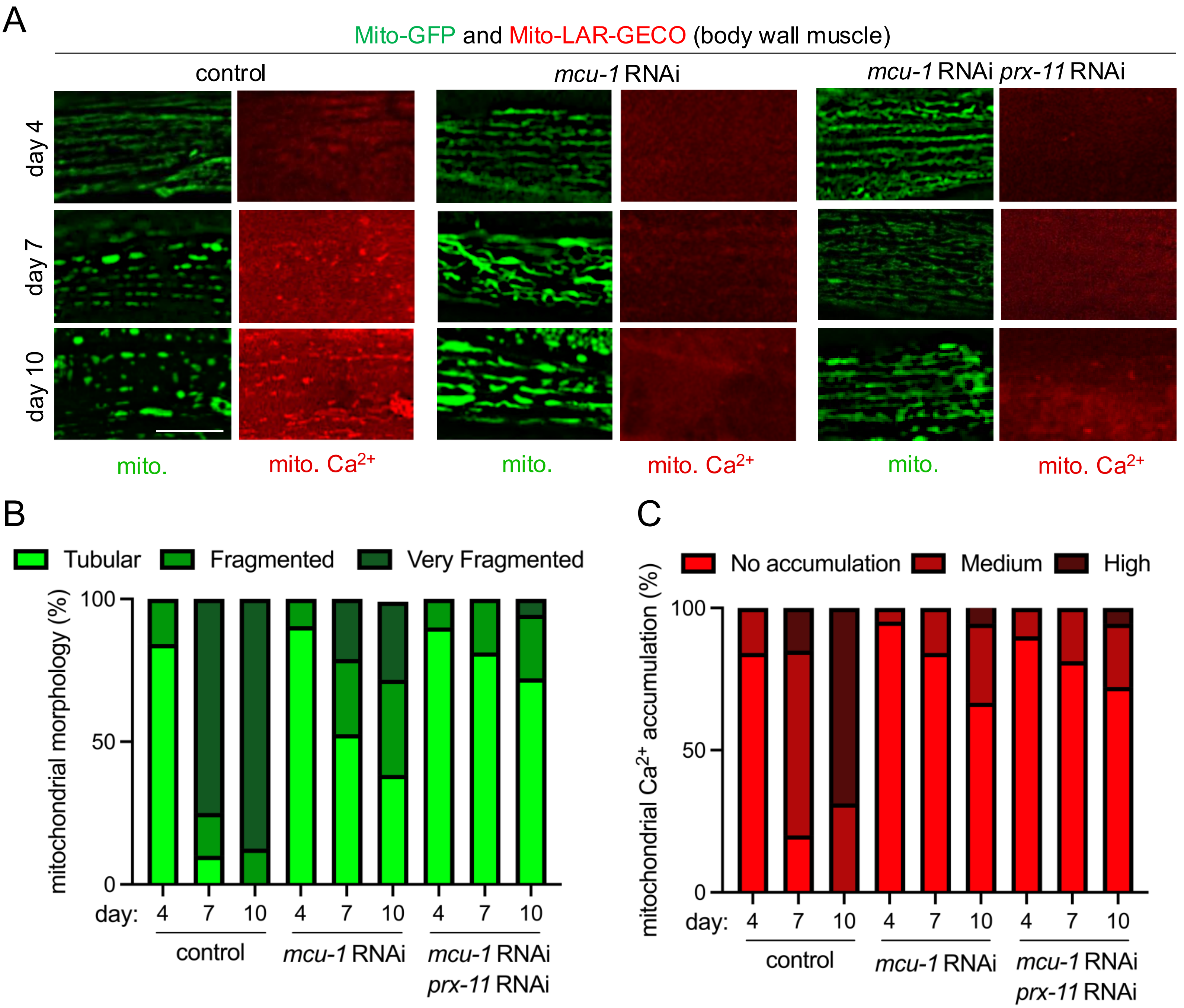

**Supplementary Figure 2. *mcu-1* knockdown suppresses age-dependent mitochondrial $Ca^{2+}$ accumulation and, partly, mitochondrial fragmentation.** (A) Representative images of Mito-GFP and the Mito-LAR-GECO $Ca^{2+}$ reporter in body wall muscle of day 4, day 7, and day 10 adult hermaphrodite animals fed either control RNAi alone, *mcu-1* RNAi alone, or *mcu-1* RNAi in combination with *prx-11* RNAi. (B,C) Classification of mitochondrial morphology and mitochondrial $Ca^{2+}$ accumulation in day 4, day 7, and day 10 adult hermaphrodite animals fed either control RNAi alone, *mcu-1* RNAi alone, or *mcu-1* RNAi in combination with *prx-11* RNAi. Bar: 10 µm.

Supplementary Figure 3

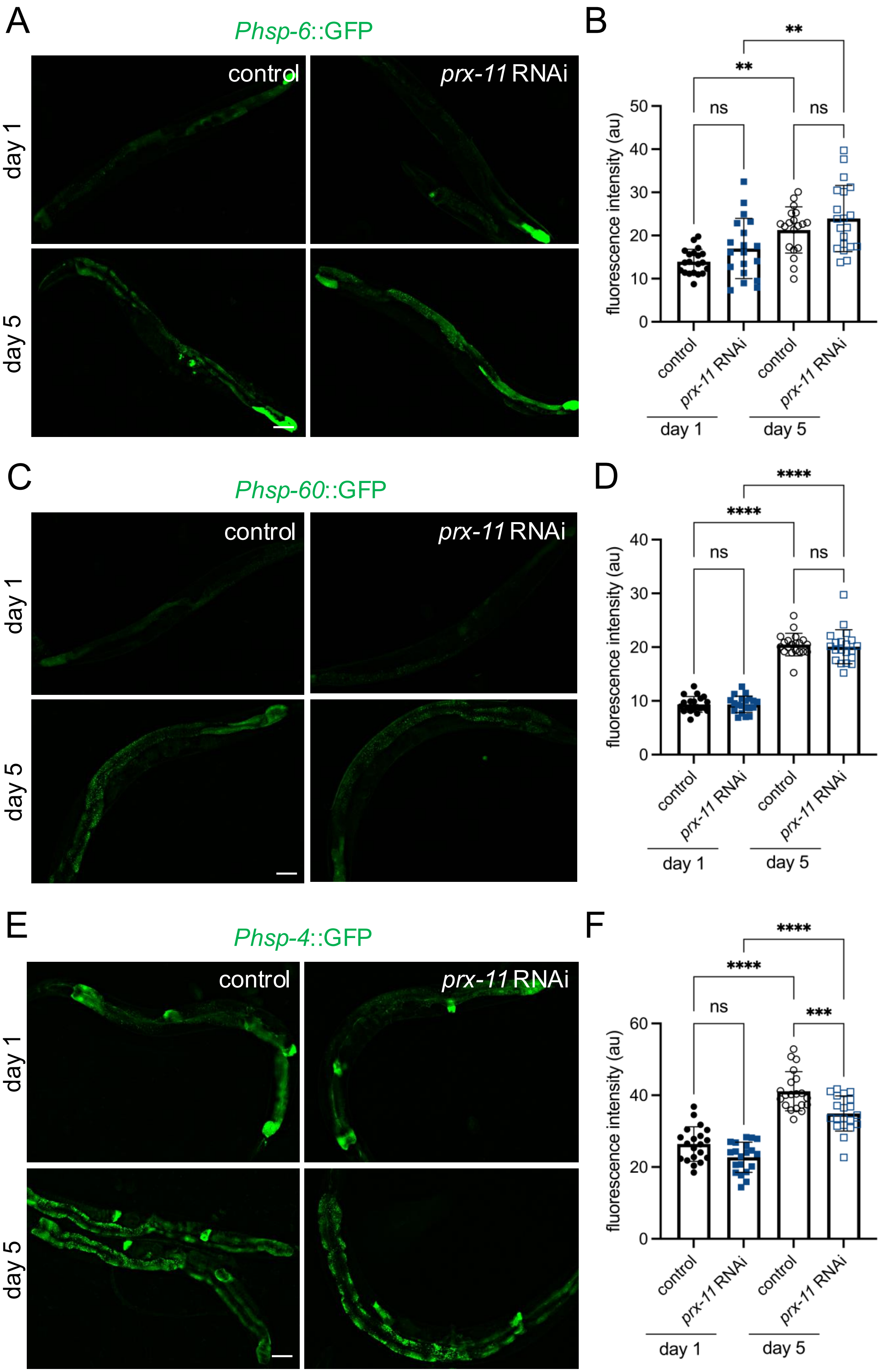

**Supplementary Figure 3. *prx-11* RNAi does not activate mitochondrial or endoplasmic reticulum unfolded protein responses.** (A) Representative images of *Phsp-6*::GFP in day 1 and day 5 adult hermaphrodite animals fed either control or *prx-11* RNAi. (B) Quantification of *Phsp-6*::GFP fluorescence intensities in day 1 and day 5 adult hermaphrodite animals fed either control or *prx-11* RNAi ($n$ = 20 worms per condition). Data are presented as mean ± SD. **, $p \leq 0.01$; ns, not significant. One-way ANOVA with Tukey's multiple comparisons. (C) Representative images of *Phsp-60*::GFP in day 1 and day 5 adult hermaphrodite animals fed either control or *prx-11* RNAi. (D) Quantification of *Phsp-60*::GFP fluorescence intensities in day 1 and day 5 adult hermaphrodite animals fed either control or *prx-11* RNAi ($n$ = 20 worms per condition). Data are presented as mean ± SD. ****, $p \leq 0.0001$; ns, not significant. One-way ANOVA with Tukey's multiple comparisons. (E) Representative images of *Phsp-4*::GFP in day 1 and day 5 adult hermaphrodite animals fed either control or *prx-11* RNAi. (F) Quantification of *Phsp-4*::GFP fluorescence intensities in day 1 and day 5 adult hermaphrodite animals fed either control or *prx-11* RNAi ($n$ = 20 worms per condition). Data are presented as mean ± SD. ****, $p \leq 0.0001$; ***, $p \leq 0.001$; ns, not significant. One-way ANOVA with Tukey's multiple comparisons. Bars: 100 μm.

Supplementary Figure 4

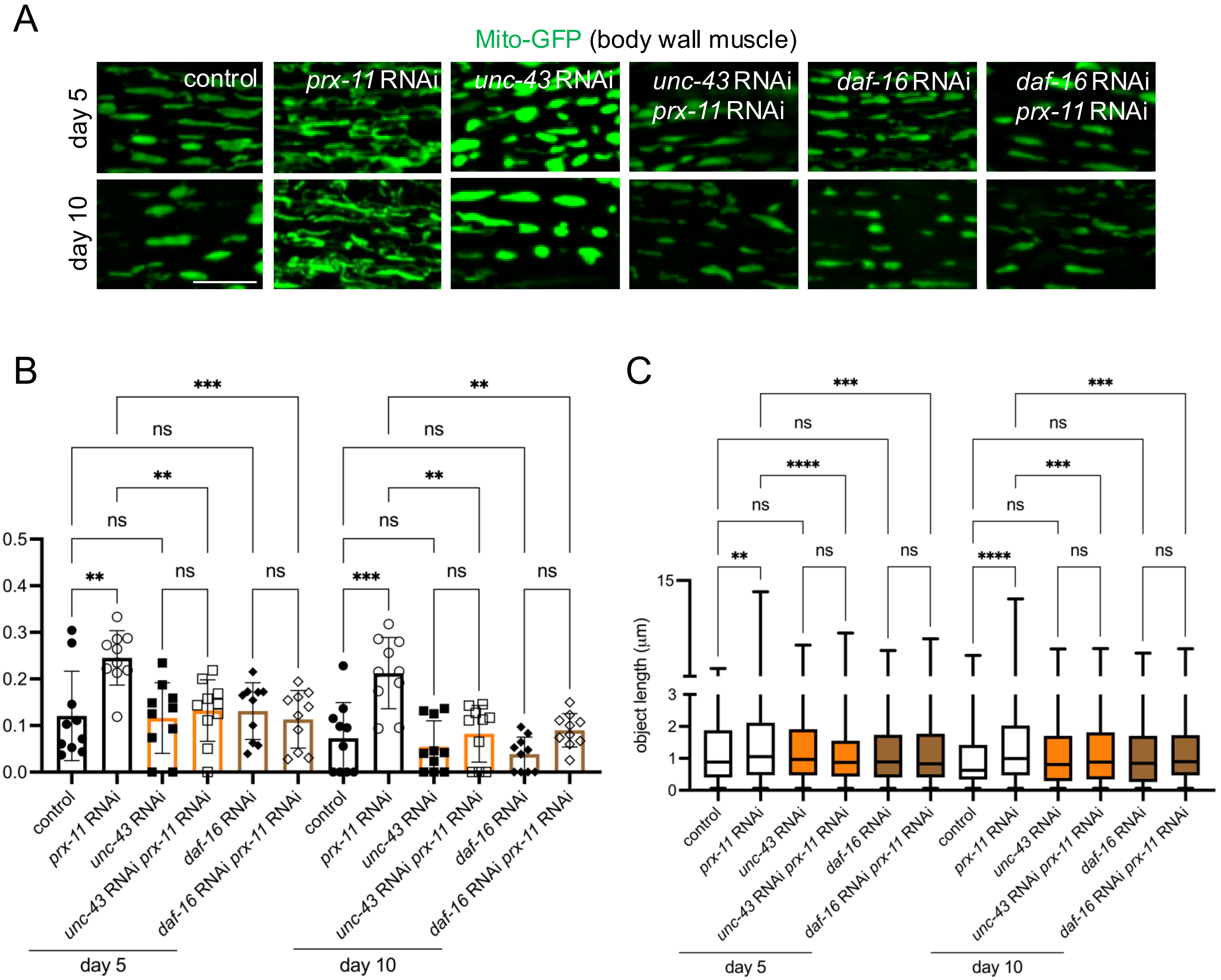

**Supplementary Figure 4. Mitochondrial tubulation during aging in *prx-11*-RNAi animals requires *daf-16* and *unc-43*.** (A) Representative images of Mito-GFP in body wall muscles of day 5 and day 10 adult hermaphrodite animals fed either control RNAi alone, *prx-11* RNAi alone, *unc-43* RNAi alone, *unc-43* and *prx-11* RNAi in combination, *daf-16* RNAi alone, or *daf-16* and *prx-11* RNAi in combination. (B) Quantification of mitochondrial junctions per object in day 5 and day 10 adult hermaphrodite animals fed either control RNAi alone, *prx-11* RNAi alone, *unc-43* RNAi alone, *unc-43* and *prx-11* RNAi in combination, *daf-16* RNAi alone, or *daf-16* and *prx-11* RNAi in combination ($n$ = 10 worms per condition). Data are presented as mean ± SD. ***, $p \leq 0.001$; **, $p \leq 0.01$; ns, not significant. One-way ANOVA with Tukey's multiple comparisons. (C) Mito-GFP object lengths in day 5 and day 10 adult hermaphrodite animals fed either control RNAi alone, *prx-11* RNAi alone, *unc-43* RNAi alone, *unc-43* and *prx-11* RNAi in combination, *daf-16* RNAi alone, or *daf-16* and *prx-11* RNAi in combination ($n$ = 10 worms per condition). Data are presented as box-and-whisker plots (minimum, 25$^{th}$ percentile, median, 75$^{th}$ percentile, maximum). ****, $p \leq 0.0001$; ***, $p \leq 0.001$; **, $p \leq 0.01$; ns, not significant. One-way ANOVA with Tukey's multiple comparisons. Bar: 10 μm.

**Supplementary Table 1. *C. elegans* strains used in this study**

| Strain Name | Genotype | Source |
|---|---|---|
| N2 | Wild-type | Lab stock |
| ATU3301 | *ccIs4251(Pmyo-3::GFP::LacZ::NLS + Pmyo-3::Mito-GFP + dpy-20(+))* I*; aceIs1(Pmyo-3::Mito-LAR-GECO + Pmyo-2::RFP)* | CGC |
| CB408 | *unc-43(e408)* IV | CGC |
| CF1038 | *daf-16(mu86)* I | CGC |
| CF1553 | *muIs84(Psod-3::GFP + rol-6(su1006))* | CGC |
| CU5991 | *fzo-1(tm1133)* II | CGC |
| PD4251 | *ccIs4251(Pmyo-3::GFP::LacZ::NLS + Pmyo-3::Mito-GFP + dpy-20(+))* I | CGC |
| SJ4005 | *zcIs4(Phsp-4::GFP)* V | CGC |
| SJ4058 | *zcIs9(Phsp-60::GFP + lin-15(+))* V | CGC |
| SJ4100 | *zcIs13(Phsp-6::GFP + lin-15(+))* V | CGC |
| | *prx-11(CSR1)* I | Huanhu Zhu Lab; Li et al., *Cell Reports* 2022 |
| KAB34 | *louIs2(Pges-1::mCherry-GFP-SKL::unc-54 UTR)* IV | Dolese et al., *Autophagy* 2022 |
| KAB111 | *louIs7(Pges1::sqst-1-mCherry-GFP::unc-54 UTR)* | Villalobos et al., *Nature Aging* 2023 |
| KAB296 | *ccIs4251(Pmyo-3::GFP::LacZ::NLS + Pmyo-3::Mito-GFP + dpy-20(+))* I; *fzo-1(tm1133) II* | This study |
| KAB323 | *prx-11(CSR1)* I; *louIs2(Pges-1::mCherry-GFP-SKL::unc-54 UTR)* IV | This study |
| KAB408 | *ccIs4251(Pmyo-3::GFP::LacZ::NLS + Pmyo-3::Mito-GFP + dpy-20(+)) prx-11(CSR1)* I | This study |
| KAB452 | *ccIs4251(Pmyo-3::GFP::LacZ::NLS + Pmyo-3::Mito-GFP + dpy-20(+))* I; *unc-43(e408) IV* | This study |
| KAB476 | *ccIs4251(Pmyo-3::GFP::LacZ::NLS + Pmyo-3::Mito-GFP + dpy-20(+)) daf-16(mu86) I* | This study |
